# Turnover rates of human muscle proteins in vivo reported in fractional, mole and absolute units

**DOI:** 10.1101/2024.01.21.576451

**Authors:** Ben N. Stansfield, Jennifer S. Barrett, Samuel Bennett, Connor A. Stead, Jamie Pugh, Sam O. Shepherd, Juliette A. Strauss, Julien Louis, Graeme L. Close, Paulo J. Lisboa, Jatin G. Burniston

## Abstract

Protein fractional turnover rates (FTR) represent measurements of flux through a protein pool, i.e. net abundance (ABD) of the protein. If protein abundance is not measured or is different between experimental conditions the interpretation of FTR data may be confounded. This project investigates the consequences of reporting turnover rates of human muscle proteins *in vivo* in mole and absolute units (that incorporate protein abundance data) compared to fractional (%/d) data that ignore protein abundance. Three physically active males (21 ± 1 years) were recruited and underwent a 12-d protocol of daily deuterium oxide (D_2_O) consumption and biopsies of vastus lateralis on days 8 and 12. Protein abundances were normalised to yeast alcohol dehydrogenase, added during sample preparation, and FTR was calculated from time-dependent changes in peptide mass isotopomer profiles. FTR and abundance data (fmol/ μg protein) were combined to calculate mole turnover rates (MTR; fmol/ μg protein/ d) and absolute turnover rates (ATR; ng/ μg protein/ d). Abundance data were collected for 1,772 proteins and FTR data were calculated from 3,944 peptides representing 935 proteins (average 3 peptides per protein). The median (M), lower- (Q1) and upper-quartile (Q3) values for protein FTR (%/d) were M = 4.3, Q1 = 2.52, Q3 = 7.84. Our analyses suggest MTR data is preferred over FTR, particularly for studies on multiprotein complexes, wherein MTR takes account of potential differences amongst the molecular weight of the component subunits. ATR data may be preferred over MTR and FTR, particularly when comparing samples with different abundance profiles.

## Introduction

In healthy adults, skeletal muscle is the largest tissue (by mass) in the human body, representing approximately 40 % of total body mass and accounting for 50 – 70 % of total body protein. Muscle proteins exist in a continuous cycle of synthesis and degradation (collectively known as protein turnover), which is essential to maintain muscle quality and facilitate changes in protein abundance profiles that underpin cellular adaptation. Losses in muscle mass are associated with heightened disease mortality [1], and changes to the turnover of muscle protein may also be an important factor underpinning muscle health. Muscle is an accessible tissue in humans and emerging techniques for dynamic proteome profiling have the potential to offer new insight into the mechanisms underpinning human muscle adaptation [2].

The dynamic nature of proteins was established in the early 20^th^ Century in work led by Rudolf Schoenheimer [3], which used a stable isotope-labelled amino acid (^15^N-tyrosine) to achieve biosynthetic labelling of newly synthesised proteins in rats *in vivo*. Muscle protein turnover has since been studied extensively but, for the most part, human data are constrained to reports on the average synthesis rate of mixed-protein samples based on the analysis of amino acid hydrolysates [4]. Soon after the turn of the century, advances in proteomic methods enabled studies on the turnover of large numbers of individual proteins, and were first conducted using ^2^H_10_-leucine in yeast [5]. The development of stable isotope labelling of amino acids in culture (SILAC) in mammalian cells [6] enabled the turnover of individual proteins in human cell cultures using ^2^H_3_-leucine. Doherty *et al* [7] reports the degradation rates of almost 600 proteins in human A549 adenocarcinoma cells using dynamic SILAC with contemporary ^13^C_6_-labelled lysine and arginine. Similarly, Cambridge *et al* [8] used dynamic SILAC in the mouse C2C12 muscle cell line and reported a median protein degradation rate of 1.6 %/h (equating to a half-life of ∼43 h) amongst 3528 proteins studied.

Dynamic SILAC requires extensive isotope labelling of amino acid precursors, which is readily achieved in cell culture but impractical in humans. Moreover, cell studies are unable to capture the complexity of human tissues *in vivo* and it is challenging to predict protein turnover rates *in vivo* from data generated in cell cultures [9]. Proteome dynamic studies in humans *in vivo* have investigated the turnover of individual proteins using the stable isotope, deuterium oxide (D_2_O or ‘heavy water’), which can be administered via a participant’s drinking water. Low levels of D_2_O consumption are safe and enable studies to be conducted under free-living conditions for periods of several days or more. Combined with peptide mass spectrometry (MS) D_2_O labelling can be employed to measure the turnover rates of individual proteins [10] and early studies in humans reported the turnover rates of specific proteins in blood [11, 12].

To our knowledge, just 5 studies [2, 13–17] report dynamic proteome data in human muscle using D_2_O, and mostly these works have investigated muscle responses to exercise training. In the majority, existing data on protein-specific turnover in human skeletal muscle *in vivo* calculate fractional synthesis rates (FSR) and report the data in percent per day (%/d) units. If protein abundance is known or assumed to be constant during the period of biosynthetic labelling, the term, fractional turnover rate (FTR), is preferred rather than FSR. Amongst the previous literature, Scalzo *et al* [13] reports the greatest number (n=381) of proteins analysed and found deuterium incorporation was greater in male compared to female participants during a 4-weeks of sprint interval training, but numerical data on the FSR of each protein was not reported. Shankaran *et al* [14] and Murphy *et al* [15] also provide protein-specific data on the FSR of 273 and 190 proteins, respectively, in human muscle but did not investigate the abundance of these proteins. Three earlier reports [2, 16, 17] from our laboratory include protein abundance (ABD) data alongside the measurements of protein-specific FSR in human muscle but the abundance and FSR data were not combined to report data in mole or absolute units.

Fractional turnover measurements provide information on the flux through the protein pool but are ignorant to the size of the pool (i.e. net protein abundance) and the potential inter-relationships amongst proteins. Muscle is renowned for its plasticity and can exhibit a broad repertoire of phenotypes, underpinned by different protein abundance profiles. For example, proteomic studies have highlighted robust changes to the abundance profile of proteins in human muscle in the contexts of exercise [18], ageing [19, 20] or disease [21]. Therefore, the abundance profile of muscle proteins may need to be considered alongside protein-specific synthesis data, particularly when comparisons are made across different populations with different muscle phenotypes. Label-free proteomics data can be normalised to spike-in standards [22] or calculated from endogenous proteins [23]. Herein, we have applied spike-in methods to investigate the abundance and turnover rates of human muscle proteins in vivo using mole (e.g. fmol/ μg total protein/ day) and absolute (e.g. ng/ μg total protein/ day) units, and we find the different unis of measurement alter the biological interpretation of protein-specific turnover data.

## Methods

### Participants

Three physically active males (21 ± 1 years; height 178 ± 1 cm; weight 75 ± 5 kg) were recruited and received verbal and written information, including potential risks, prior to providing written informed consent. The study was approved by the Liverpool John Moores School of Sport and Exercise Science Research Committee (M18SPS006) and conformed with the Declaration of Helsinki, except registration in a publicly accessible database.

### Experimental Protocol

Figure 1 provides an overview of the experimental protocol which consisted of a 12-day cross-sectional, observation study including metabolic labelling of newly synthesised protein *in vivo* using deuterium oxide (^2^H_2_O; D_2_O) administration. Seventy-two hours prior to the labelling period, preparatory data including anthropological measurements and peak aerobic capacity, were collected. Baseline saliva, blood, and muscle samples were collected on day 0, prior to daily D_2_O administration. Saliva and venous blood samples were collected on days 0, 4, 8 and 12 to measure body water deuterium enrichment. Muscle samples were also collected on days 8 and 12 via micro-needle biopsy of the vastus lateralis.

**Figure 1.**
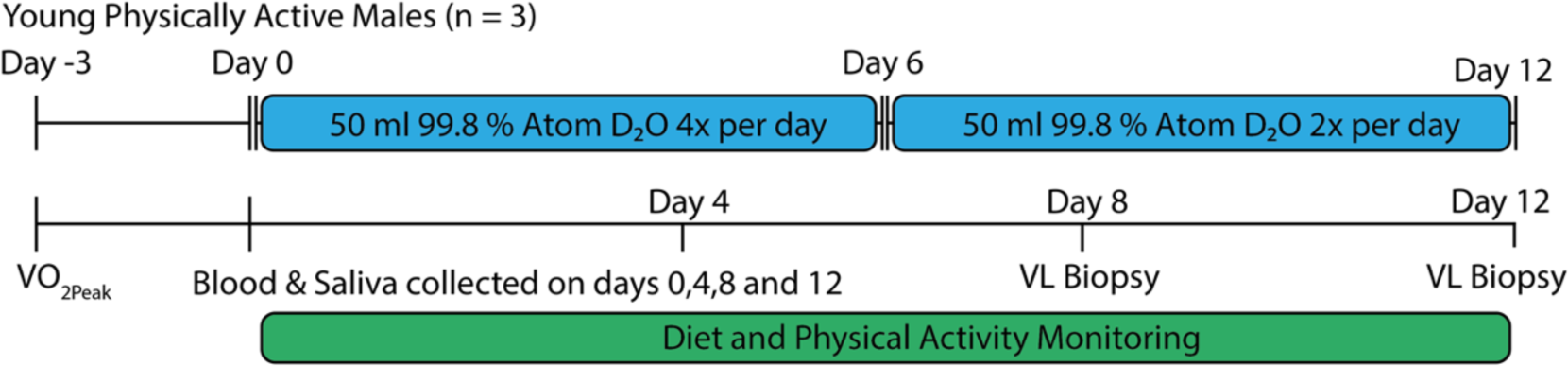
Experiment design. Visual schematic of the experimental design employed in the current study. Performance measures and biometric data (outlined in Table 1) were collected 3 days prior to the study commencement. Blood and saliva were collected on days 0, 4, 8 and 12, to calculate D_2_O precursor enrichment. Fifty millilitres of 99.8 % D_2_O was consumed by all participants 4x per day for the first 6 days. This dose was lowered to 2x per day for days 6 – 12. Vastus lateralis biopsies were taken on day 8 and day 12 of the experimental period. Diet and physical activity monitoring were conducted throughout the experimental period (day 0-12).

**Table 1.**
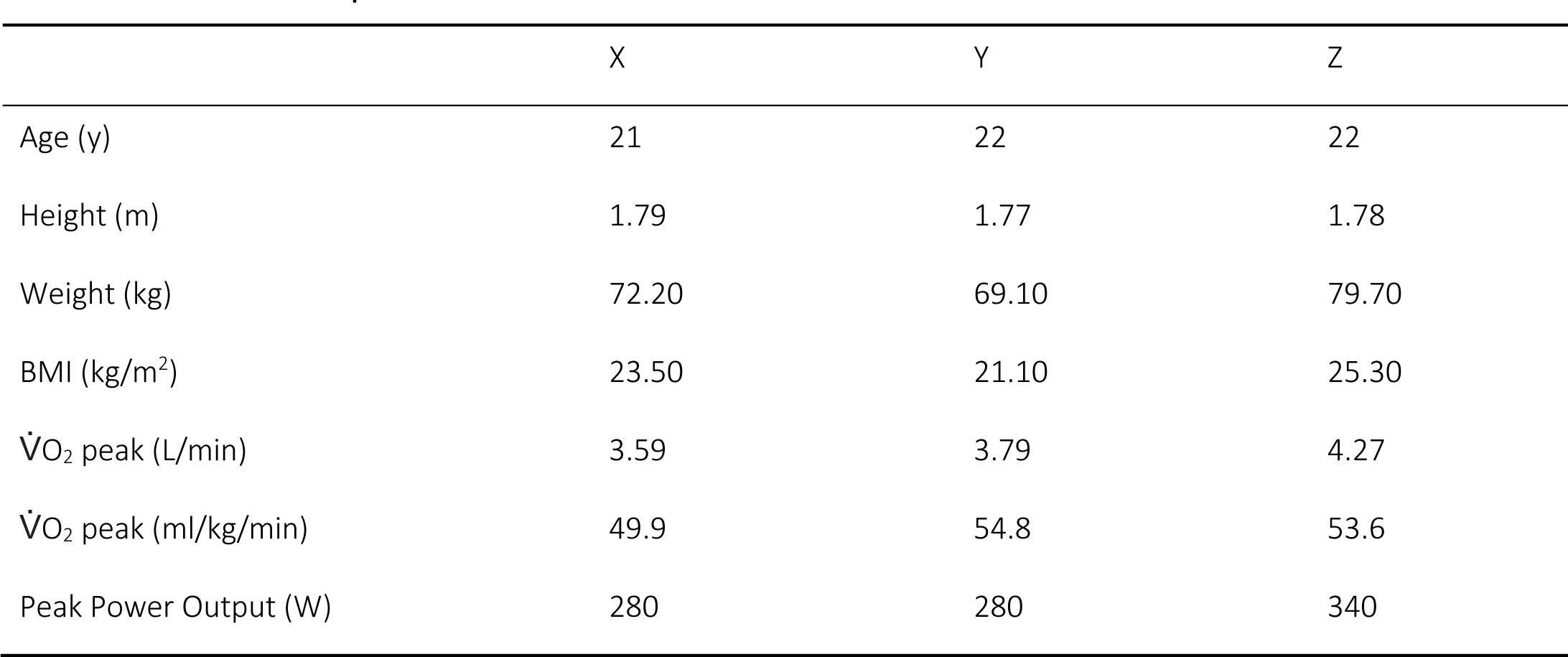
Participant characteristics.

|  | X | Y | Z |
| --- | --- | --- | --- |
| Age (y) | 21 | 22 | 22 |
| Height (m) | 1.79 | 1.77 | 1.78 |
| Weight (kg) | 72.20 | 69.10 | 79.70 |
| BMI (kg/m<sup>2</sup>) | 23.50 | 21.10 | 25.30 |
| $\dot{V}O_2$ peak (L/min) | 3.59 | 3.79 | 4.27 |
| $\dot{V}O_2$ peak (ml/kg/min) | 49.9 | 54.8 | 53.6 |
| Peak Power Output (W) | 280 | 280 | 340 |

### Assessment of aerobic exercise capacity

Participants attended the laboratory in the morning after an overnight fast and their resting heart rate and blood pressure were measured (DINAMAP V100, General Healthcare, UK) in a seated position. Peak oxygen uptake (VrO_2_ peak) was measured, using an incremental exercise test to volitional exhaustion on a cycle ergometer (Lode, Groningen, The Netherlands). Respiratory gases were measured using an online gas collection system (CORTEX Biophysik MetaLyzer 3B stationary CPX). The test consisted of an initial load of 100 W for 10 minutes, followed by 30 W increases in external load at 2-minute intervals until cadence reduced to <50 rpm, at which point the test was terminated. VrO_2_ peak was reported as the mean oxygen uptake during the final 1-minute of exercise.

### Stable isotope labelling *in vivo*

Participants were instructed to maintain their habitual exercise and dietary routine throughout the 12-day D_2_O labelling period. Participants recorded the duration and rate of perceived exertion (RPE) of each training session using TrainingPeaks software (TrainingPeaks, Denver, CO, USA). Dietary intake was monitored by recording meal information using a smartphone application (MyFitnessPal, Under Armour, Baltimore, MD, USA) [24].

Biosynthetic labelling of newly synthesised proteins was achieved by oral consumption of deuterium oxide (D_2_O). Consistent with our previous work [2, 16], participants consumed 50 ml of 99.8 atom % of D_2_O four times per day for days 1-5 commencing after the first muscle biopsy on day 0. On days 6 -12, the dosage was lowered to 50 ml two times per day. All 50 ml doses were dispensed in a nutrition laboratory and sealed prior to distribution to the participants. Participants were instructed to consume each dose ∼4 hours apart to negate any potential side effects e.g., nausea.

Saliva samples were collected, upon waking, by each participant using pre-labelled saliva collection kits (Salivette, Sarstedt, NC, USA). Participants delivered saliva samples to the laboratory at each visit using a cooled container. Collection tubes were centrifuged for 2 mins at 1000 x *g* and aliquots of saliva were stored at -80 °C until analysis.

Venous blood samples were collected in EDTA-coated vacutainer tubes via a single use butterfly needle (Beckton Dickson, UK) inserted into the antecubital fossa. Plasma was extracted by centrifugation (1,200 x *g*, 4 °C for 10 min) prior to storage at -80 °C for subsequent analysis.

### Muscle Biopsy Protocol

Muscle samples were obtained from the vastus lateralis of the participant’s dominant leg after an overnight fast using a Bard Monopty Disposable Core Biopsy Instrument 12-gauge x 10 cm length (Bard Biopsy System, Tempe, AZ). Local anaesthesia (0.5 % Marcaine) was administered, and a 0.5 cm longitudinal incision was made through the skin. The muscle fascia was then pierced, and 2-3 muscle pieces were taken to collect adequate amounts (minimum 50 mg) of sample. Samples were blotted to remove excess blood, and visible fat and connective tissue were removed through dissection. Muscle tissue was snap-frozen in liquid nitrogen and stored at -80 °C for subsequent analysis.

### Calculation of D_2_O Enrichment

Body water enrichment of D_2_O was measured in plasma and saliva samples against external standards that were constructed by adding D_2_O to PBS over the range from 0.0 to 5.0 % in 0.5 % increments. Deuterium enrichment of aqueous solutions was determined by gas chromatography-mass spectrometry after exchange to acetone [25]. Samples were centrifuged at 12,000 g, 4 °C for 10 min, and 20 µl of sample supernatant or standard was reacted overnight at room temperature with 2 µl of 10 M NaOH and 4 µl of 5 % (v/v) acetone in acetonitrile. Acetone was then extracted into 500 µl chloroform and water was captured in 0.5 g Na_2_SO_4_ before transferring a 200-µl aliquot of chloroform to an auto-sampler vial. Samples and standards were analysed in triplicate by using an Agilent 5973 N mass selective detector coupled to an Agilent 6890 gas chromatography system (Agilent Technologies, Santa Clara, CA, USA). A CD624-GC column (30 m, 30.25 mm^3^ 1.40 mm) was used in all analyses.

Samples (1 μl) were injected by using an Agilent 7683 autosampler. The temperature program began at 50 °C and increased by 30 °C/min to 150 °C and was held for 1 min. The split ratio was 50:1 with a helium flow of 1.5 ml/min. Acetone eluted at ∼3 min. The mass spectrometer was operated in the electron impact mode (70 eV) and selective ion monitoring of m/z 58 and 59 were performed by using a dwell time of 10 ms/ ion.

### Muscle processing

Muscle samples were pulverized in liquid nitrogen, then homogenized on ice in 10 volumes of 1 % Triton X-100, 50 mM Tris, pH 7.4 (including complete protease inhibitor; Roche Diagnostics, Lewes, United Kingdom) using a PolyTron homogenizer. Homogenates were incubated on ice for 15 min, then centrifuged at 1000 x *g*, 4 °C, for 5 min to fractionate insoluble (myofibrillar) proteins from soluble proteins. Soluble proteins were decanted and cleared by further centrifugation (12,000 x *g*, 4 °C, for 45 min). Insoluble proteins were resuspended in a half-volume of homogenization buffer followed by centrifugation at 1000 x *g*, 4 °C, for 5 min. The washed pellet was then solubilized in lysis buffer (7 M urea, 2 M thiourea, 4 % CHAPS, 30 mM Tris, pH 8.5) and cleared by centrifugation at 12,000 x *g*, 4 °C, for 45 min. Protein concentrations of the insoluble and soluble protein fractions were measured by Bradford assay. Aliquots containing 500 µg protein were precipitated in 5 volumes of ice-cold acetone and incubated for 1 h at -20 °C. Proteins were then resuspended in lysis buffer to a final concentration of 5 μg/ μl.

Tryptic digestion was performed using the filter-aided sample preparation (FASP) method [26]. Aliquots containing 100 µg protein were precipitated in acetone and resuspended in 40 μl UA buffer (8 M urea, 100 mM Tris, pH 8.5). Samples were transferred to filter tubes and washed with 200 µl of UA buffer. Proteins were incubated at 37 °C for 15 min in UA buffer containing 100 mM dithiothreitol followed by incubation (20 min at 4 °C) protected from light in UA buffer containing 50 mM iodoacetamide. UA buffer was exchanged with 50 mM ammonium bicarbonate and sequencing-grade trypsin (Promega, Madison, WI, USA) was added at an enzyme to protein ratio of 1:50. Digestion was allowed to proceed at 37 °C overnight then peptides were collected in 100 μl 50 mM ammonium bicarbonate containing 0.2 % trifluoroacetic acid. Samples containing 4 µg of peptides were de-salted using C_18_ Zip-tips (Millipore) and resuspended in 20 µl of 2.5 % (v/v) ACN, 0.1 % (v/v) formic acid (FA) containing 10 fmol/ μl yeast alcohol dehydrogenase (ADH1; MassPrep, Waters Corp., Milford, MA).

### Liquid chromatography-mass spectrometry of the myofibrillar protein fraction

Liquid chromatography-mass spectrometry of myofibrillar proteins was performed using nanoscale reverse-phase ultra-performance liquid chromatography (NanoAcquity; Waters Corp., Milford, MA) and online electrospray ionization quadrupole-time-of-flight mass spectrometry (Q-TOF Premier; Waters Corp.). Samples (5 μl corresponding to 1 μg tryptic peptides) were loaded by partial-loop injection on to a 180 μm ID x 20 mm long 100 Å, 5 µm BEH C_18_ Symmetry trap column (Waters Corp.) at a flow rate of 5 μl/ min for 3 min in 2.5 % (v/v) ACN, 0.1% (v/v) FA. Separation was conducted at 35 °C via a 75 μm ID x 250 mm long 130 Å, 1.7 µm BEH C_18_ analytical reverse-phase column (Waters Corp.). Peptides were eluted using a non-linear gradient that rose to 37.5 % acetonitrile 0.1% (v/v) FA over 90 min at a flow rate of 300 nl/ min. Eluted peptides were sprayed directly into the mass spectrometer via a NanoLock Spray source and Picotip emitter (New Objective, Woburn, MA). Additionally, a LockMass reference (100 fmol/ μl Glu-1-fibrinopeptide B) was delivered to the NanoLock Spray source of the mass spectrometer at a flow rate of 1.5 μl/ min and was sampled at 240 s intervals. For all measurements, the mass spectrometer was operated in positive electrospray ionization mode at a resolution of 10,000 full width at half maximum (FWHM). Before analysis, the time-of-flight analyser was calibrated using fragment ions of [Glu-1]-fibrinopeptide B from m/z 50 to 1990.

Mass spectra for liquid chromatography-mass spectrometry profiling were recorded between 350 and 1600 m/z using mass spectrometry survey scans of 0.45-s duration with an interscan delay of 0.05 s. In addition, equivalent data-dependent tandem mass spectra (MS/MS) were collected from each baseline (day 0) sample. MS/MS spectra of collision-induced dissociation fragment ions were recorded from the 5 most abundant precursor ions of charge 2+ 3+ or 4+ detected in each survey scan. Precursor fragmentation was achieved by collision-induced dissociation at an elevated (20–40 eV) collision energy over a duration of 0.25 s per parent ion with an interscan delay of 0.05 s over 50–2000 m/z. Acquisition was switched from MS to MS/MS mode when the base peak intensity exceeded a threshold of 30 counts/s and returned to the MS mode when the total ion chromatogram (TIC) in the MS/MS channel exceeded 50,000 counts/s or when 1.0 s (5 scans) were acquired. To avoid repeated selection of peptides for MS/MS, the program used a 30-s dynamic exclusion window.

### Liquid chromatography-mass spectrometry of the myofibrillar protein fraction

Data-dependent label-free analysis of soluble protein fractions was performed using an Ultimate 3000 RSLC nanosystem (Thermo Scientific, Waltham, MA) coupled to a Fusion mass spectrometer (Thermo Scientific). Samples (3 μl corresponding to 600 ng of protein) were loaded on to the trapping column (Thermo Scientific, PepMap100, C_18_, 75 μm X 20 mm), using partial loop injection, for 7 minutes at a flow rate of 9 μl/min with 0.1 % (v/v) trifluoroacetic acid. Samples were resolved on a 500 mm analytical column (Easy-Spray C_18_ 75 μm, 2 μm column) using a gradient of 96.2 % A (0.1 % formic acid) 3.8 % B (79.9 % ACN, 20 % water, 0.1 % formic acid) to 50 % B over 90 min at a flow rate of 300 nL/min. The data-dependent program used for data acquisition consisted of a 120,000-resolution full-scan MS scan (AGC set to 4^e5^ ions with a maximum fill time of 50 ms) and MS/MS using quadrupole ion selection with a 1.6 m/z window, HCD fragmentation and normalised collision energy of 32 and LTQ analysis using the rapid scan setting and a maximum fill time of 35 msec. The machine was set to perform as many MS/MS scans as to maintain a cycle time of 0.6 sec. To avoid repeated selection of peptides for MS/MS the program used a 60 s dynamic exclusion window.

### Label-free quantitation of protein abundances

Progenesis Quantitative Informatics for proteomics (Non-Linear Dynamics, Newcastle, UK) was used to perform label-free quantitation on samples collected at days 8 and 12 only. QToF data were LockMass corrected using the doubly charged monoisotopic ion (*m/z* 785.8426) of the Glu-1-fibrinopeptide B. Prominent ion features were used as vectors to warp each data set to a common reference chromatogram and an analysis window of 15–105 min and 350–1500 m/z was selected. Log-transformed MS data were normalized by inter-sample abundance ratio, and relative protein abundances were calculated using unique peptides only. Abundance data were normalised to the median abundance of 3 most abundant peptides of yeast ADH1 [22] to derive abundance measurements in fmol/μg units. MS/MS spectra were exported in Mascot generic format and searched against the Swiss-Prot database (2018.7) restricted to Homo-sapiens (20,272 sequences) using a locally implemented Mascot server (v.2.2.03; www.matrixscience.com). Enzyme specificity was trypsin, allowing 1 missed cleavage, carbamidomethyl modification of cysteine (fixed). QToF data was searched using m/z errors of 0.3 Da, whereas FUSION data were searched using MS errors of 10 ppm and MS/MS errors of 0.6 Da. Mascot output files (xml format), restricted to nonhomologous protein identifications, were recombined with MS profile data in Progenesis.

### Measurement of protein turnover rates

Mass isotopomer abundance data from samples collected at days 8 and 12 were extracted from MS spectra using Progenesis Quantitative Informatics (Non-Linear Dynamics, Newcastle, UK). Consistent with our previous work, e.g. [27], the abundances of the monoisotopic peak (m_0_), m_1_, m_2_ and m_3_ mass isotopomers were collected over the entire chromatographic peak for each proteotypic peptide that was used for label-free quantitation of protein abundances. Mass isotopomer information was processed in R version 3.5.2 (R core team., 2016). Incorporation of D_2_O into newly synthesized protein results in a decrease in the molar fraction of the monoisotopic (*f*m_0_) peak that follows the pattern of an exponential decay. The rate constant (*k)* for the decay of *f*m_0_ was calculated as a first-order exponential spanning from the beginning (*t0*) (day 8) to end (*t*) (day 12) of the D_2_O labelling period (Equation 1).

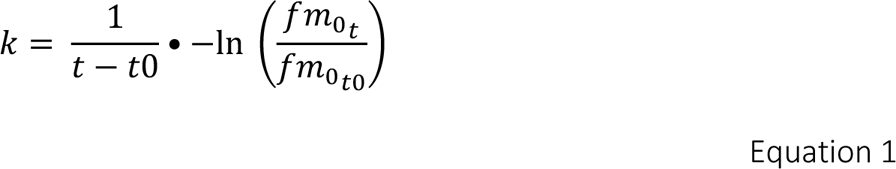

The rate of change in mass isotopomer distribution is also a function of the number of exchangeable H-D sites, and this was accounted for by referencing each peptide sequence against standard tables reporting the relative enrichment of amino acids by deuterium in humans [11] to give the fractional turnover rate (FTR) for each peptide. Individual protein FTR was reported as the median of peptide values assigned to each protein (decimal values were multiplied by 100 to give FTR in %/d) for each participant. Mole turnover rate (MTR, fmol/μg/d) was calculated by multiplying protein FTR (expressed as a decimal) by the mole abundance of the protein normalised to the yeast ADH1 spike-in. Absolute turnover rate (ATR, ng/μg/d) was calculated by multiplying MTR by the predicted molecular weight (kDa) of each protein. Protein half-life (t_1/2_) in days was estimated from decimal FTR data by Equation 2.

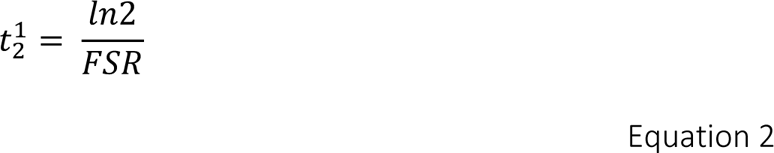

### Bioinformatic Analysis

Functional annotation and the association of proteins with pathways of the Kyoto Encyclopaedia of Genes and Genomes [KEGG; http://www.genome.jp/kegg/, [28]] were conducted using the Perseus platform [29]. Protein interactions were investigated using bibliometric mining in the Search Tool for the Retrieval of Interacting Genes/proteins (STRING; http://string-db.org/) [30]. Protein physio-chemical characteristics, including isoelectric point (*pI*) and molecular weight (MW) were calculated using the Swiss Institute of Bioinformatics EXpasy ProtParam tool (https://web.expasy.org/protparam/). Statistical analysis was performed in R (Version 3.6.2). Within-subject differences between samples collected on day 8 and day 12 were investigated by repeated measures one-way ANOVA. Significance was identified as *P* ≤ 0.05 and a false-discovery rate of 5 % calculated from q-values [31]. Differences amongst myosin heavy chain (MyHC) isoforms and the subunits of multiprotein complexes were investigated by one-way ANOVA with Tukey’s HSD post-hoc analysis. Pearson’s moment correlation analyses were used to determine relationships between protein abundance and turnover rate expressed in relative (e.g., FTR or t_1/2_), mole and absolute units. Amino acid sequence logos were generated using Seq2Logo 2.0 (https://services.healthtech.dtu.dk/services/Seq2Logo-2.0/). Stoichiometry of multiprotein complexes were calculated and compared to the expected stoichiometry reported within The Complex Portal [32]. Gene ontology analysis was conducted on the top quantile of proteins when ranked by ABD, FTR, MTR and ATR using cluster profiler with significance set at an adjusted P value ≤ 0.05.

## Results

### Protein abundance measurements

Three physically active age-matched males were studied that had similar peak aerobic capacity and body mass index (Table 1). In total, 1,885 proteins were identified and 1,772 of these proteins had at least 1 unique peptide (<1 % FDR) detected in both day 8 and day 12 samples in all 3 participants. The abundance of muscle proteins spanned 6 orders (Figure 2A) of magnitude from 0.003 fmol/μg (leucine tRNA Ligase; SYLC) to 4146.35 fmol/μg (haemoglobin subunit beta; HBB). There were no statistically significant differences in protein abundance between day 8 and day 12 and the R^2^ for protein abundance data was 0.987 (Figure 2B). The median coefficient of variation in protein abundance was 6 % with an inter-quartile range from 2.97 % to 12.54 %, which demonstrates a high level of repeatability between protein abundance measurements at day 8 and day 12. Gene ontology analysis of the upper quartile (n = 234 proteins) of protein abundances, included proteins associated with muscle contraction, muscle system process, regulation of muscle contraction, sarcomere organization and striated muscle contraction as the most enriched processes in human muscle (Figure 2C). Whereas biological processes, including tRNA aminoacylation, positive regulation of transcription, protein folding, cell migration and regulation of translation were enriched amongst proteins in the lower quartile of abundance measurements.

**Figure 2.**
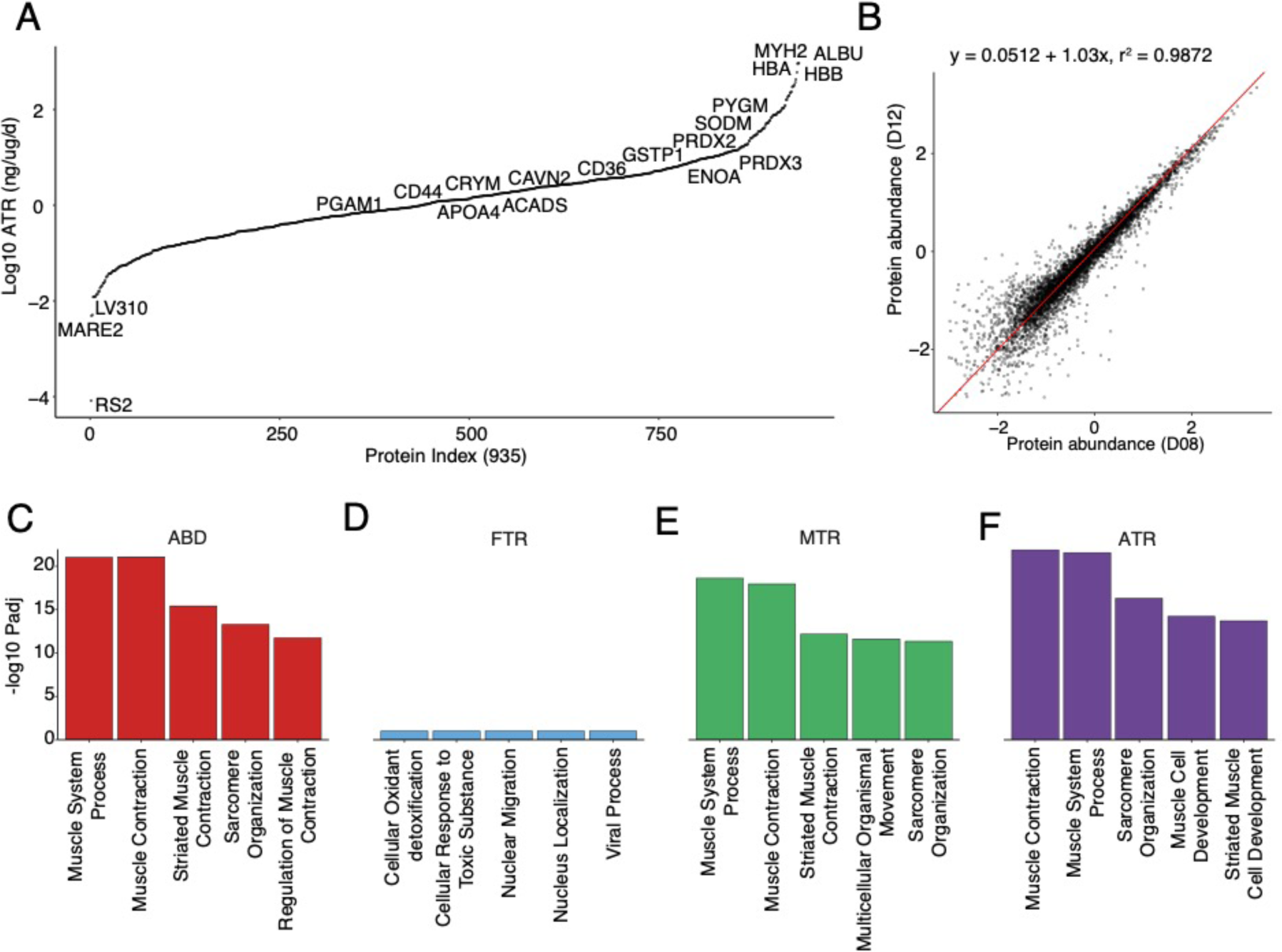
Proteome profiling of human vastus lateralis muscle in vivo. Proteins were extracted from the vastus lateralis of humans and turnover rates for 935 proteins were quantified in at least one participant. (A) Log10 transformed distribution plot, of proteins ranked by ATR (ng/μg/d). Proteins of interest are labelled using their UniProt ID. (B) Linear regression of within-subject protein abundance data at day 8 and day 12. Panels C-F report the 5 most significant enriched GO Biological Processes amongst the top-ranking proteins contained within the upper quartile when the protein dataset was ranked by either (C) abundance, (D) fractional, (E) molar or (F) absolute turnover rate.

### Protein turnover measurements

Body water enrichment measured in blood plasma (1.71 ± 0.08 %) on day 8 was not different (P = 0.1058) from values (1.89 ± 0.07 %) measured on day 12. Stringent filters were applied to select peptides with clearly resolved envelopes of m_0_, m_1_, m_2_ and m_3_ mass isotopomers, and in all 6,800 protein-specific peptides met the inclusion criteria for turnover calculations in one or more participants. The turnover rates of 935 proteins were measured in at least 1 participant (Suppl Table S1), whereas data were collected for 766 proteins in 2 or more participants and the synthesis rate of 444 proteins was measured in all 3 participants. Unless otherwise stated, data are presented from at least n = 2 participants in the subsequent text and figures. The turnover of individual proteins in human vastus lateralis *in vivo* ranged from 0.32 %/d (microtubule associated protein RP/EB family member 2; MARE2) to 54.43 %/d (nuclear protein localisation protein 4 homolog; NPL4) and the median (IQR) of protein-specific FTR was 4.3 (2.52 – 7.84) %/d. MTR had a median of 0.04 (IQR: 0.01 – 0.10) fmol/μg/d and ranged between 0.00013 (MARE2) and 56.89 fmol/μg/d haemoglobin subunit beta (HBB). ATR values had a median of 1.53 (IQR: 0.48 – 4.63) ng/μg/d and ranged between 0.005 (MARE2) and 931.98 ng/μg/d (albumin; ALBU). Different gene ontological classifications were highlighted amongst the 75^th^ percentile of proteins when turnover data were presented in relative or absolute units (Figure 2). When ranked by FTR, cellular oxidant detoxification, cellular response to toxic substance, nuclear migration, nucleus localization and viral process were amongst the top-ranked terms. (Figure 2D). In contrast, proteins ranked in the 75^th^ percentile by MTR were associated with terms, multicellular organismal movement, muscle contraction, muscle system process, sarcomere organization, and striated muscle contraction (Figure 2E). And proteins ranked in the 75^th^ percentile by ATR were associated with the terms muscle cell development, muscle contraction, muscle system process, sarcomere organization and striated muscle cell development (Figure 2F). Furthermore, antioxidant enzymes, including SODM (mitochondrial superoxide dismutase [Mn]), PRDX3 (peroxiredoxin 3), PRDX2 (peroxiredoxin 2) and (GSTP1) glutathione S-transferase-P were also amongst the top-ranked proteins by ATR, consistent with their prominent role in skeletal muscle physiology.

Our analysis encompassed 35 myofibrillar proteins, including each of the main components of the sarcomere (Figure 3), which were primarily extracted from the insoluble fraction. The myosin heavy chain (MyHC) isoform profile was 65 ± 2.4 % type IIa (MYH2), 24 ± 3.5 % type I (MYH7) and 12 ± 1.2 % type IIx (MYH1). Accordingly, the abundance (ABD) of MYH2 (634.50 ± 173.79 fmol/ μg) was significantly (p ≤ 0.01) greater than MYH1 (114.67 ± 38.25 fmol/μg) and MYH7 (225.97 ± 24.96 fmol/μg), whereas there were no significant differences amongst turnover of MyHC isoforms (Figure 3B) when expressed in FTR (%/d) units. When rate data were expressed in mole (Figure 3C) or absolute terms (Figure 3D), the turnover profile of MyHC isoforms more closely mirrored the abundance data and both the MTR (8.07 ± 1.93 fmol/μg/d) and ATR (1799.2 ± 429.82 ng/μg/d) of MYH2 were significantly (p ≤ 0.01) greater than MYH7 (MTR; 2.56 ± 1.13 fmol/μg/d, ATR; 571 ± 252.3 ng/μg/d) and MYH1 (MTR; 2.03 ± 1.04 fmol/μg/d, ATR; 453.70 ± 232.19 ng/μg/d).

**Figure 3.**
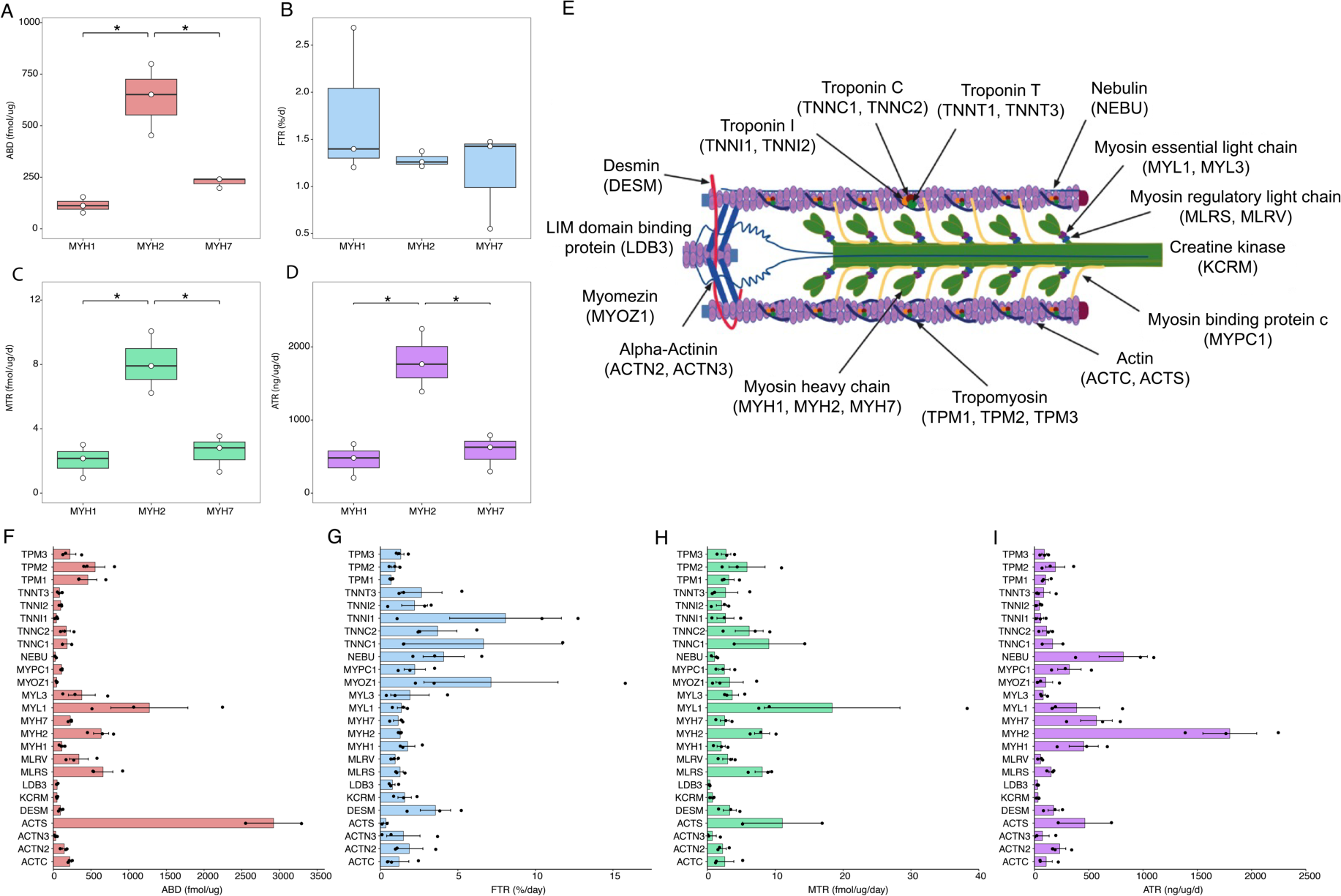
Turnover rates of muscle sarcomeric proteins. Box plots representing the MyHC profile of the vastus lateralis are displayed for abundance (A), FTR (B), MTR (C) and ATR (D) measurements (N = 3). *Significant (P ≤ 0.05) differences between proteins determined by ANOVA and TUKEY’s HSD. Panel E is a visual representation of the skeletal muscle sarcomere with major proteins labelled. Protein abundance (F), FTR (G), MTR (H) and ATR (I) measurements for sarcomeric proteins detected in the insoluble fraction (N = 2-3) are presented as individual points. Bars represent mean values ± SE. All proteins are labelled using their UniProt ID.

Protein turnover data were collected for 48 proteins of the major energy metabolism pathways in human muscle (Figure 4), including fatty acid β-oxidation (17 of 42 annotated in gene ontology databases), glycolysis (15 of 67) and the TCA cycle (16 of 30). The median (IQR) ABD of proteins involved in fatty acid oxidation and the TCA cycle were 2.11 (IQR; 1.13 – 3.65) fmol/μg and 3.34 (IQR; 0.63 – 8.41) fmol/μg respectively, whereas the ABD for proteins involved in glycolysis was approximately 10-fold greater, 37.00 (IQR; 9.12 – 105.56) fmol/μg. Despite the markedly greater abundance of glycolytic enzymes, the median FTR (2.10, 1.20 – 2.40 %/d) of glycolytic proteins was not different from enzymes of either TCA cycle (2.20, 1.66 – 3.04 %/d) or fatty acid oxidation pathway (2.78, 2.19 – 3.94 %/d). Differences amongst the turnover of proteins of the different metabolic pathways were more transparent when data were expressed in mole or absolute units. The MTR (0.69, 0.15 – 1.34 fmol/μg/d) and ATR (29.23, 7.84 – 81.48 ng/μg/d) of enzymes involved in glycolytic processes was ≥10-fold greater than the TCA cycle (MTR; 0.077, 0.022 – 0.134 fmol/μg/d, ATR; 4.06, 1.24 – 9.41 ng/μg/d) and fatty acid oxidation (MTR; 0.065, 0.039 – 0.095 fmol/μg/d, ATR; 2.12, 1.29 – 3.63 ng/μg/d).

**Figure 4.**
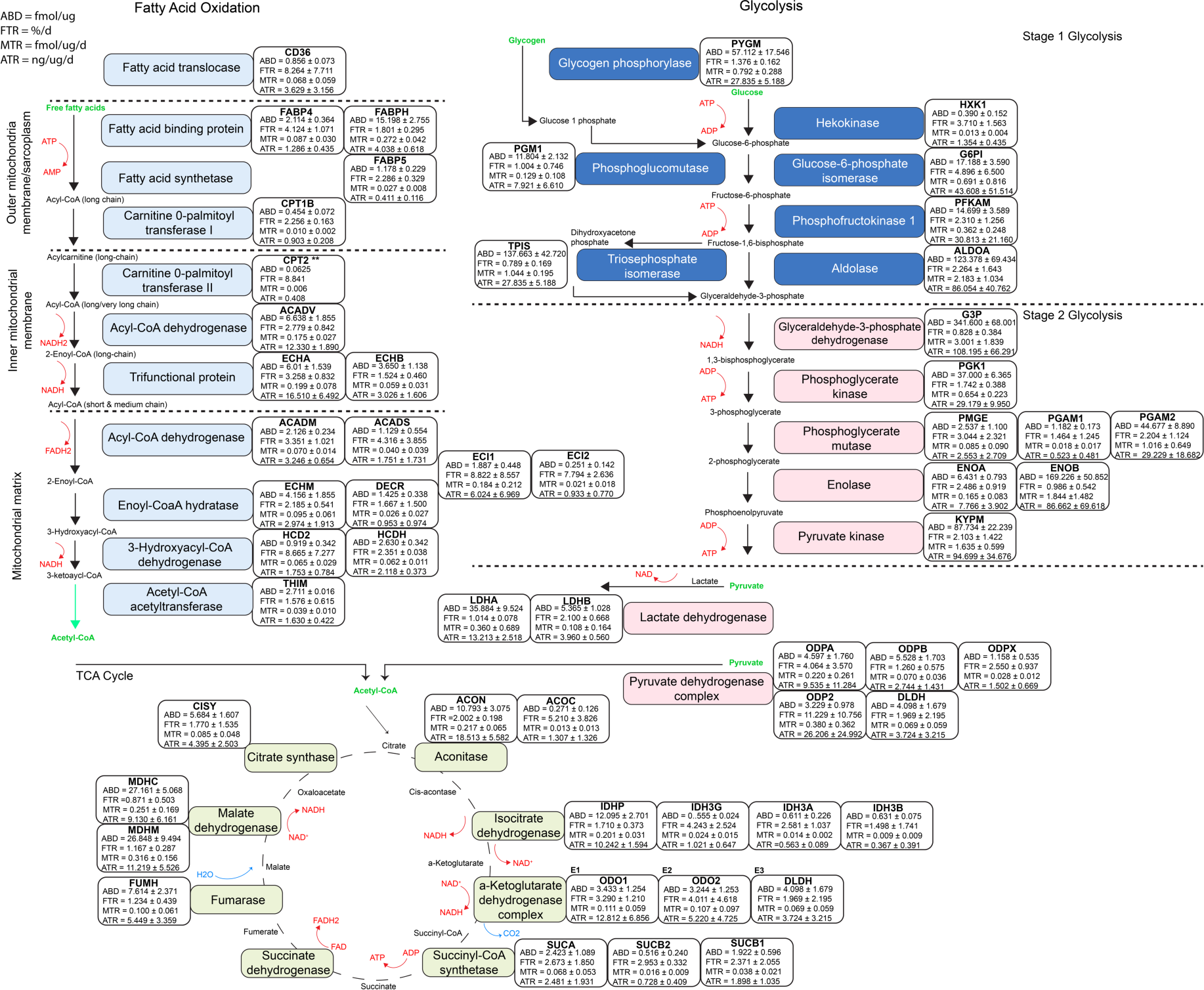
Proteomic profiling of metabolic enzyme pathways. Individual protein abundance (ABD) turnover rates in fractional (FTR), mole (MTR) and absolute (ATR) units for proteins involved in fatty acid oxidation, glycolysis and the TCA cycle are reported Data are mean ± SD from *n* = 2-3 participants except for CPT2, which was measure in 1 participant only. Common names of proteins are labelled in coloured boxes, with adjacent boxes labelled using UniProt protein IDs.

### Predictors of protein turnover rates in human muscle in vivo

Protein turnover expressed in fractional units (FTR, %/d) did not correlate (r = -0.082) with mole protein abundance (Figure 5A), whereas MTR exhibited a strong (r = 0.9695) positive relationship (Figure 5B) and ATR somewhat correlated with mole protein abundance (r = 0.6964) (Figure 5C). Neither protein molecular weight (MW; Figure 5 D-F) nor isoelectric point (p*I*; Figure 5 G-I) corelated with protein turnover expressed in either relative or absolute units. The median (IQR) for p*I* and MW of proteins in the current study were 41.44 (IQR; 25.97 – 63.45) KDa and pH 6.105 (IQR; 5.37 – 7.58). Pearson’s corelation analysis of the top 50 proteins with the lowest and highest t_1/2_ values found no correlation (r = -0.0024) with predicted protein t_1/2_ values calculated using the N-end rule of degradation. Similarly, there was no significant enrichment of linear motifs amongst either the top (Figure 5K) or bottom-ranked proteins (Figure 5L).

**Figure 5.**
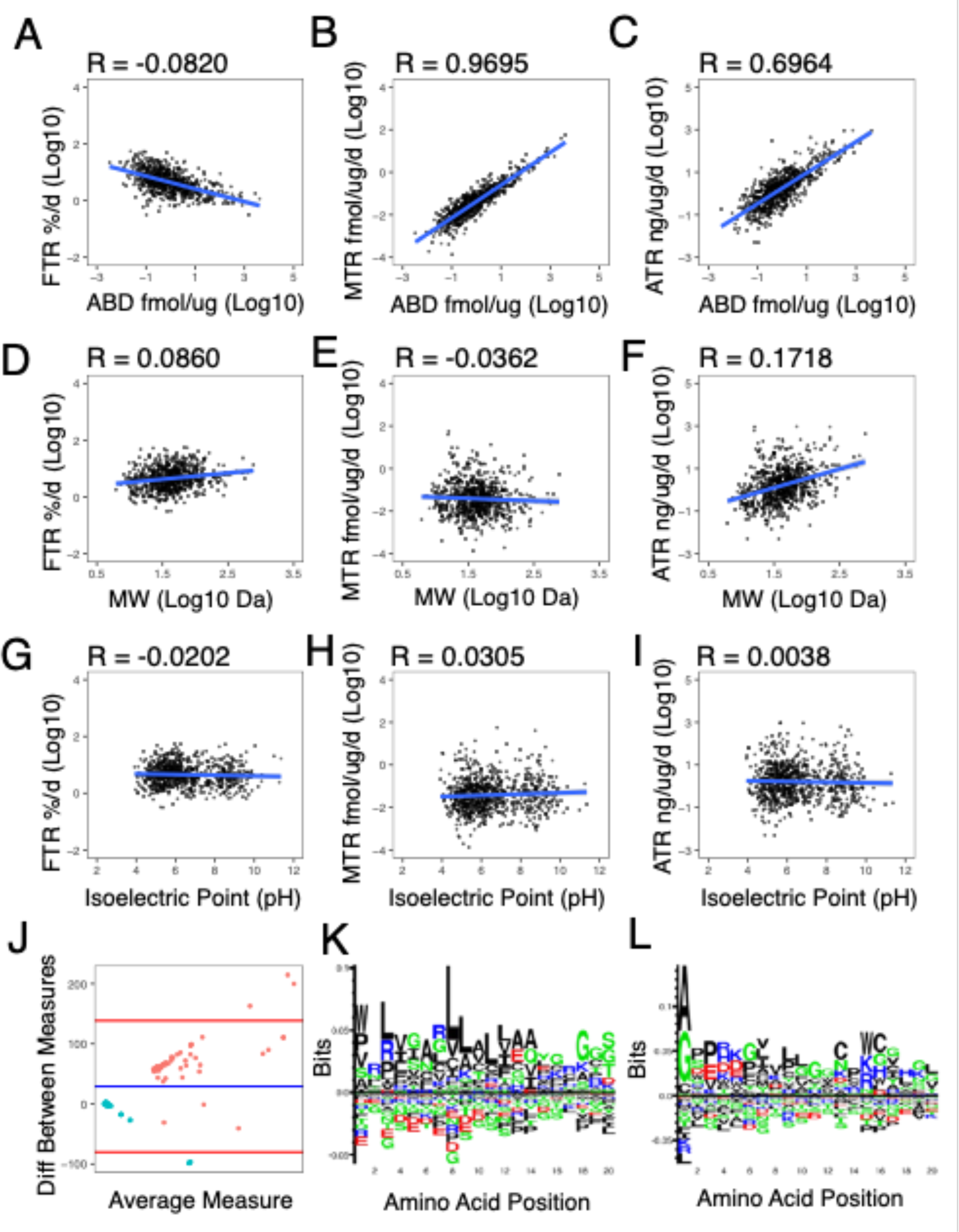
Predictors of Protein Turnover Rates. (A-I) Each turnover measurement was correlated (Pearson’s) with protein abundance, MW, and P*I* to attempt to predict the turnover of individual proteins. (J) Bland Altman plot of the top 50 proteins with the shortest (blue) and longest (red) protein t_1/2_ values against the predicted protein t_1/2_ values calculated using the N end rule of degradation. (K-L) P weighted Kullback-Leibler logos of the top quartile (K) and bottom (L) quartile of proteins ranked by t_1/2_.

### Subunit stoichiometry of multiprotein complexes

Data were collected for 79 subunits of 6 multiprotein complexes (MPC), including the 26S proteasome and complexes I, II, III, IV and V of the mitochondrial respiratory chain (Figure 6). On average the mean ± SD mole abundance of the 26S proteasome (0.342 ± 0.224 fmol/μg) was significantly less (P ≤ 0.05) than the abundance of respiratory chain complexes III, IV and V (Figure 6A). Subunits of Complex I (1.414 ± 0.856 fmol/ug) were significantly less abundant than subunits of Complex III (7.311 ± 4.453 fmol/μg) and Complex V (11.335 ± 14.349 fmol/ug). Similarly, subunits of Complex II (2.328 ± 1.944 fmol/ug) and complex IV (4.833 ± 3.835 fmol/ug) were also significantly less abundant than Complex V. There were significant (P ≤ 0.05) differences in FTR (%/d) between the 26S proteasome and mitochondrial respiratory chain complexes III and V (Figure 6B). In contrast to ABD data the turnover of the 26S Proteasome subunit (8.436 ± 7.03450 %/day) was significantly higher than that of Complex III (1.946045 ± 0.479 %/day) and Complex V (2.210 ± 1.091 %/day). Turnover data expressed in mole (Figure 6C) and absolute terms (Figure 6D) were in better agreement with protein abundance measures, for example Complex IV (0.21 ± 0.25 fmol/μg/day) and Complex V (0.180 ± 0.181 fmol/ug/day) had significantly (P ≤ 0.05) greater mean ± SD MTR than the 26S proteasome (0.027 ± 0.029 fmol/μg/day). Complex IV also had a significantly higher MTR than complex I (0.09 ± 0.12 fmol/μg/day). Absolute values of MPC highlighted the ATR of Complex V was significantly (P ≤ 0.05) greater (6.83 ± 11.62 ng/μg/day) than complex I (2.46 ± 3.66 ng/μg/day) and the 26S proteasome (1.02 ± 0.98 ng/μg/day).

**Figure 6.**
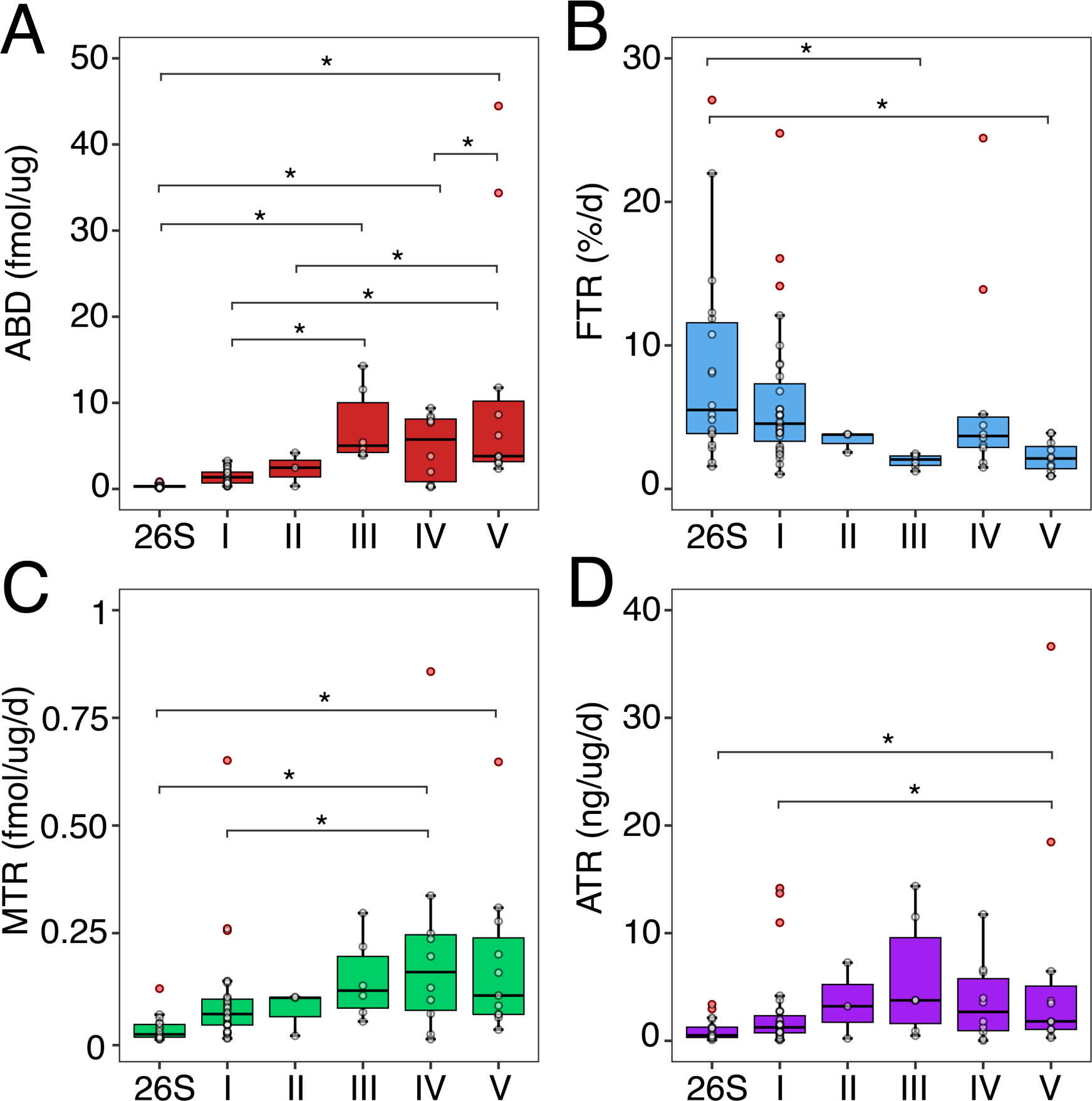
Proteomic profiling of multiprotein complexes in humans in vivo. Representative box plots of the ABD (A), FTR (B), MTR (C) and ATR (D) for proteins identified within each of the mitochondrial respiratory chain complexes (I-V) and the 26S proteasome. Individual dots represent mean values (n = 2-3 participants) for each protein within each multiprotein complex (MPC). Outliers are highlighted in red and median values for the corresponding MPC are indicated by solid black lines within the box plot. * Indicate complexes that are significantly different from one another as determined by ANOVA and Tukey’s HSD (P ≤ 0.05).

Individual subunits within a MPC display differing turnover rates. We identified 18 of the 47 subunits of the 26S proteasome in at least n = 2 participants and 24 subunits in *n* = 1 participants (Figure 7). Ten subunits belonged to the 20S core particle and 13 to the 19S regulatory particle. FTR values of subunits within the 26S proteasome ranged from 1.57 ± 1.06 %/d (proteasome subunit alpha type 3; PSA3) to 27. 09 ± 17.50 %/d (26S proteasome AAA-ATPase subunit RPT1; PRS7), with a combined median of 5.50 %/d (IQR; 3.86 – 11.58) for the 18 proteins identified in the current work. MTR and ATR values ranged from 0.003 ± 0.002 fmol/μg/d (PSA3) to 0.12 ± 0.16 fmol/μg/d (proteasome subunit alpha type 7; PSA7) and 0.09 ± 0.06 ng/μg/d (PSA3) to 3.38 ± 4.59 ng/μg/d (PSA7) respectively. The median (IQR) MTR and ATR for the 18 proteins identified in the 26S proteasome are reported as 0.015 (0.008 – 0.038) fmol/μg/d and 0.51 (0.31 – 1.27) ng/μg/d respectively.

**Figure 7.**
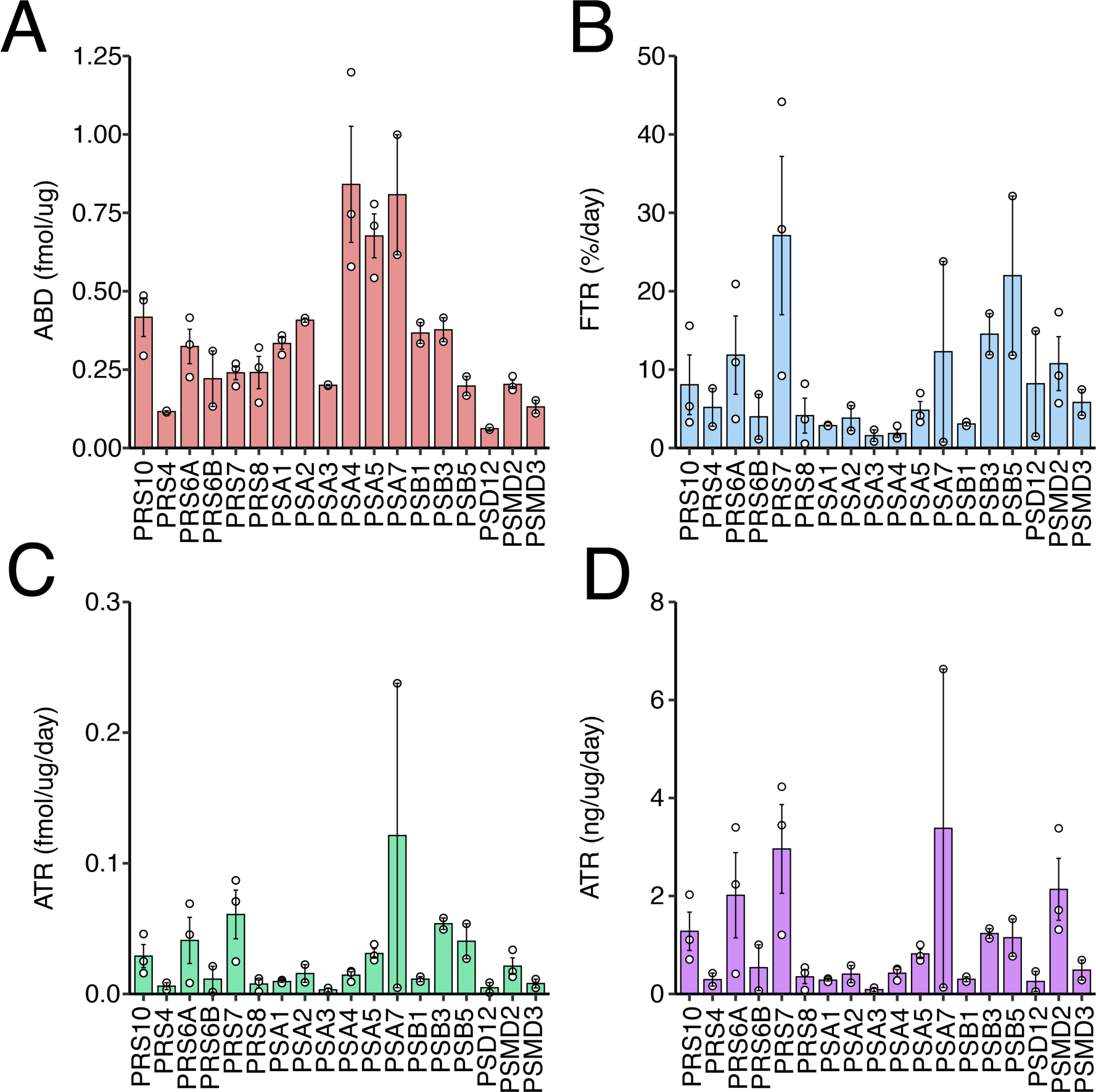
Proteomic profiling of individual subunits of the 26S Proteasome. Proteomic data were extracted for individual proteins contained within the 26S proteasome (*n* = 2-3 participants). ABD (A), FTR (B), MTR (C) and ATR (D) measurements are reported. Bars and error-bars represent mean ± SD, respectively. Points represent values for individual participants.

Complex V of the mitochondrial respiratory chain is a well-defined MPC. We herein quantified the ABD and stoichiometry of Complex V (ATP synthase) and compared the measured protein abundance of each subunit, against their predicted abundance from the Complex Portal database (Figure 8A). We also compared individual protein subunit turnover in fractional (Figure 8C), mole (Figure 8D) and absolute (Figure 8E) terms against protein abundance stoichiometry. Figure 8F-H illustrates the relationship between the turnover rates and relative protein abundance of subunits of ATP synthase. Values are expressed as a percentage of the total of ATP synthase. There was no relationship (r^2^ = - 0.144) between the FTR of a subunit and its relative protein ABD within the ATP synthase MPC (Figure 8F). For example, ATPB made up ∼ 37 % of the ABD of ATP synthase, but only 6 % of turnover (FTR). In contrast, MTR (r^2^ = 0.851) and ATR (r^2^ = 0.933) data exhibited significant relationships with the relative ABD of ATP synthase subunits (Figure 8 G-H). The measured stoichiometry of the average abundance of ATP synthase subunits when normalised to ATPG (i.e. abundance ATPA:ATPB:ATPD per fmol of ATPG) was 9.17: 11.62: 0.84 fmol/µg per fmol/µg of ATPG.

**Figure 8.**
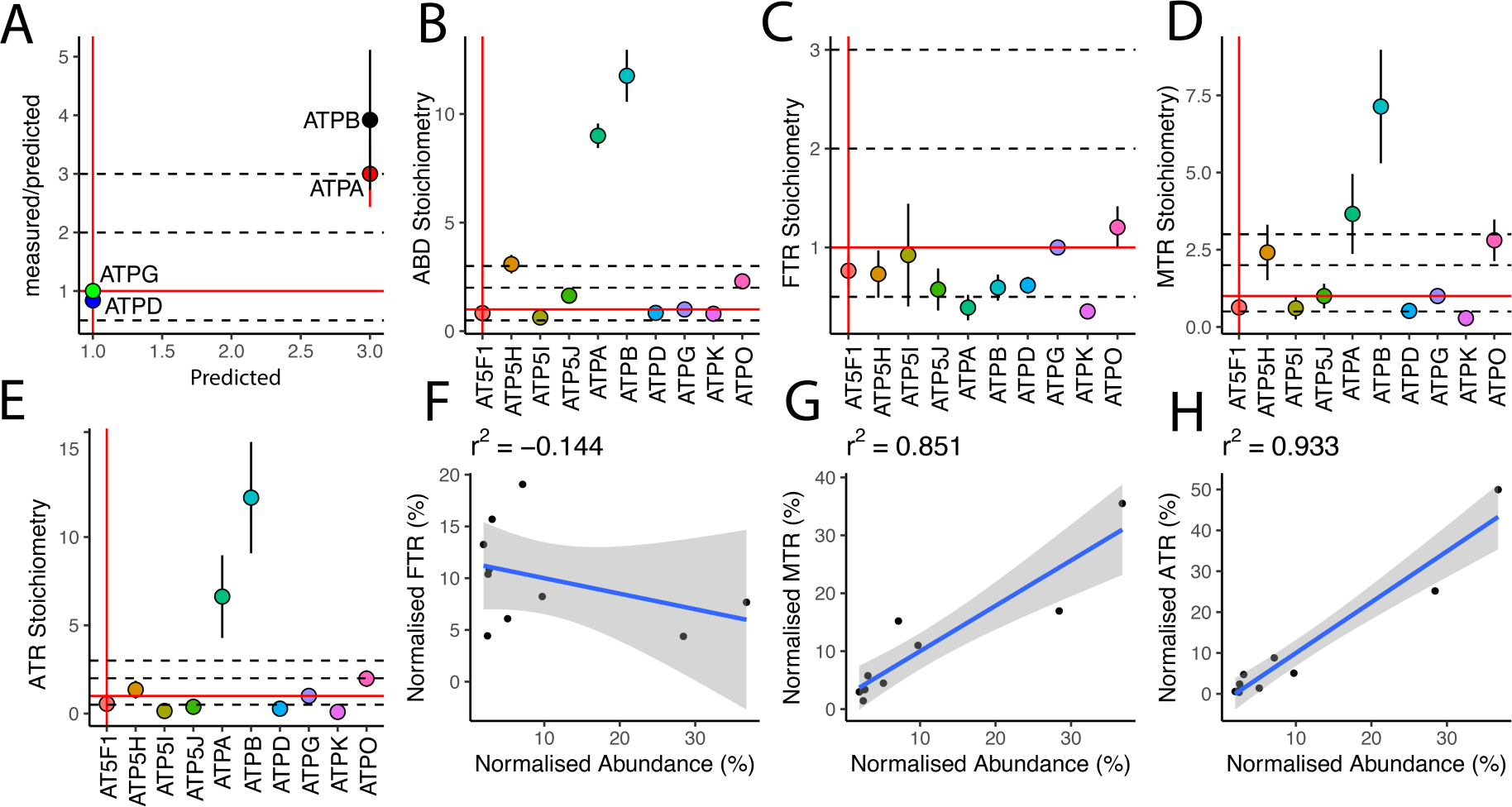
Subunit stoichiometry of mitochondrial respiratory chain Complex V - ATP Synthase, and relationship between protein turnover and protein abundance. (A) Predicted vs measured abundance values of subunits with known stochiometric ratios in humans normalized to the γ-subunit (ATPG), which forms the central stalk of ATP synthase. The stoichiometry for all quantified subunits in the data set are reported for ABD (B), FTR (C), MTR (D) and ATR (E). Protein measurements are normalised to the regulatory subunit ATPG. Measurements of ATPG are represented by red lines and dashed lines represent 0.5x 2x and 3x the reported value of ATPG. Linear regression between the abundance of subunits of ATP synthase and turnover rates in (F) fractional, (G) molar and (H) absolute units.

## Discussion

Currently few data exist on protein-specific turnover in human skeletal muscle *in vivo*, and the turnover rates of individual proteins have seldom been reported alongside protein abundance measurements. Muscle is renowned for its plasticity and can exhibit a broad range of different phenotypes underpinned by different protein abundance profiles. Therefore, it may be important to consider the abundance profile of muscle proteins alongside protein-specific turnover data. Herein, we report that the unit of measurement used for dynamic proteomic data influences the biological interpretation of the findings. Although commonplace, the presentation of protein turnover or synthesis data in fractional terms (i.e. %/d), masks the different size (molecular weight; MW) and abundance of individual proteins. Data presented in mole units that account for differences in MW amongst protein subunits may be preferred for studies on multiprotein complexes within samples or subcellular fractions. Alternatively, data in absolute units that account for differences in protein abundance between samples may be preferred for studies on longitudinal adaptation or cross-sectional analyses on differing populations.

Changes in protein abundance are underpinned by the balance between the synthesis and degradation of individual proteins [33] but this relationship does not equate to a correlation between the turnover rate of a protein and its abundance (Figure 5). Disparities between the FTR and abundance of proteins make it difficult to draw conclusions about the physiological state of muscle from FTR data alone [34]. In the current work, FTR data highlighted biological processes (Figure 2) that are not usually viewed as being prominent in healthy muscle, and provided limited insight into the allocation of cellular resources or top-ranking molecular processes. Mole turnover rates incorporate quantification of protein abundance in mole terms and provide new insight into stoichiometric relationships amongst the synthesis of subunits that form multiprotein complexes in muscle (Figure 6). Both mole and absolute synthesis rates correlated (Figure 5) with protein abundance measurements and better reflected the biological processes associated with skeletal muscle (Figure 2).

The median FTR of 935 proteins in human vastus lateralis muscle was 4.3 (IQR 2.52 – 7.84) %/d, which equates to a median protein half-life of 16 (IQR 9 – 27) days. These values appear to differ from values (∼1-2 %/d) reported in previous D2O proteomic studies [15, 16] that surveyed fewer proteins. As the depth of proteome coverage (i.e. number of proteins studied) increases it becomes increasingly important to account for the abundance and molecular weight of each protein when estimating the gross average turnover of muscle protein. Large or highly abundant proteins (e.g., myosin and actin) contribute more to the gross average turnover of protein in muscle than less abundant proteins that may exhibit relatively higher rates of turnover. For example, when our FTR data is weighted using absolute abundance values, (i.e. ATR) the pooled FTR for human vastus lateralis muscle was 1.4 %/d and in line with previous findings in active individuals. Notably, muscle proteins that exhibit high absolute turnover rates and therefore contribute the most to the gross average turnover rate, include blood proteins (e.g. albumin and haemoglobin subunits), myoglobin (MYG) and glycolytic enzymes. In total, the top 10 proteins ranked by ATR account for almost 30 % of protein turnover in human skeletal muscle. The prominent contribution of blood proteins and glycolytic enzymes sheds new light on the interpretation of earlier mixed protein FTR data generated by analysis of amino acid hydrolysates. Such analyses of muscle sarcoplasmic fractions often failed to find differences in response to ageing or diet and exercise interventions. If the small number of top-ranking ATR proteins are unaffected by the intervention this will overshadow potentially important responses by proteins that rank lower in ATR but are nonetheless biologically important.

Because FTR data do not consider the different abundances or molecular weights of the proteins studied, data expressed in MTR or ATR units may be more appropriate for investigating the stoichiometry amongst subunits of multi-protein complexes. Consistent with studies in yeast [35], we report the half-life of 26S proteasome subunits is shorter than that of most respiratory chain subunits. However, the relative abundance of respiratory chain complexes is greater than the proteasome. The mitochondrial respiratory chain comprises five multi-protein complexes and the relative combination and organisation of each complex influences mitochondrial super complex formation [36], and is crucial for subunit-complex stability [37]. The stoichiometry amongst subunits may be adaptable, for example differences in physical activity levels [20] and exercise training [38] are associated with different abundance profiles of respiratory chain protein subunits. ATP-synthase (respiratory chain Complex V) is responsible ATP resynthesis [39], and defects in ATP-synthase structure are associated with diseases and ageing [40], whereas the beta subunit of ATP-synthase is particularly responsive to exercise [41].

We quantified the ABD of individual proteins constituting the ATP-synthase complex to investigate the stoichiometry amongst its subunits and their synthesis rates in human muscle *in vivo*. Data were normalised to ATPG, which constitutes the singular central stalk (gamma subunit) of each ATP synthase multiprotein complex. A stoichiometry of 1:9:12 was found between ATPG: ATPA and ATPB abundance, which differs from human mitochondrial DNA models in yeast [42] that report ATP-synthase contains 3 subunits each of ATPA and ATPB for every 1 subunit of ATPG. The greater ratio of ATPA and ATPB subunits in human muscle *in vivo* may indicate a pool of subunits that are not assembled into mature complexes. This could either present a burden to proteostasis mechanisms, including transport systems of the inner and outer mitochondrial membranes, or represent a beneficial, ready source of replacement subunits that helps to maintain the quality of the multiprotein complex. The FTR of ATP synthase subunits was similar when normalised relative to the FTR of ATPG, but the relative relationship between subunits was different (Figure 9) when data were expressed in MTR units. While our current data can be used for within-sample comparisons of FTR, MTR and ATR data, further work is required to investigate the stoichiometry of ATP-synthase subunits using purified subcomplexes or mitochondria-enriched fractions. Notably, our current data from the soluble protein fractions, which encompasses both cytosolic and mitochondrial compartments cannot distinguish complexes from individual subunits or the potential different cellular locations of proteins.

Theoretically, the turnover rate of a protein may be dictated by both its intrinsic properties, including sequence elements and physiochemical properties, and by extrinsic factors, including the cell environment and regulatory processes acting on protein translation and degradation. In HeLa cells, highly abundant proteins have slower than average synthesis rates than low abundant proteins [43] and in our data (Figure 5A) FTR tended to be inversely related (r-0.082) to protein abundance. In yeast, the intrinsic properties of proteins, including physiochemical properties, linear sequence motifs, biological function and mRNA half-life, provide some predictive relationship with protein turnover under stable culture conditions [35]. N-terminal linear sequence motifs have been associated with high- and low-turnover rate proteins but we report no relationship between N-terminal sequence and protein half-life (Figure 5J), which is consistent with earlier studies using sequence information from protein databases [7, 43], or empirically determined sequences of N-terminal peptides using tandem mass spectrometry [44]. We also found the intrinsic properties of a protein such as their predicted molecular weight (MW) and isoelectric point (p*I*) have little to no relationship with protein turnover rates *in vivo* reported in FTR units (Figure 5), which is consistent with studies in human cells *in vitro* [7, 43]. When studied using 2-dimensional gel electrophoresis, the majority of human muscle proteins resolve as multiple proteoforms [45], which cannot be distinguished in LC-MS/MS analyses of tryptic peptide digests used here and in previous studies [7, 43]. In particular, proteoforms with different post-translational modifications may exhibit marked differences in p*I* from their predicted values and this may contribute to the lack of association between FTR and predicted p*I* in the current data. However, in rat soleus [46], proteoforms of creatine kinase share similar turnover rates, whereas the turnover rate differed between the 2 proteoforms of albumin studied.

The half-life of proteins is shorter *in vitro* compared to studies on intact tissue *in vivo* [9]. In addition, the influence of extrinsic factors on the turnover of proteins may be more prominent *in vivo* than *in vitro,* even when studied, as here, under steadystate conditions. In rats the turnover rate of a particular protein *in vivo* is not always consistent across different muscles from an individual animal [46]. For example, the FTR of the primary proteoform of the blood protein, albumin, is similar (range 4.8 %/d – 6.3 %/d) regardless of whether the data is extracted from samples of heart, diaphragm, soleus or extensor digitorum longus. In contrast, the FTR of the cardiac/ slow muscle myosin essential light chain (MYL3) is 7.4 %/d in EDL, 10.7 %/d in diaphragm and 6.4 %/d in heart [46]. Moreover, the rank order of protein turnover rates is not consistent amongst different muscles [46] and the turnover rate of a protein can change in response to environmental stimuli, including muscle contraction and exercise training [2, 33]. Therefore, attempts to predict protein turnover *in vivo* from the intrinsic properties of the protein seem unlikely to be rewarding.

## Conclusion

In summary, the units chosen for reporting protein turnover data affect the biological interpretation of dynamic proteome profiling studies. MTR data is preferred over FTR, particularly for studies on multiprotein complexes, wherein MTR takes account of potential differences amongst the molecular weight of the component subunits. ATR data may be preferred over MTR and FTR, particularly when the aim is to compare between samples that may exhibit different abundance profiles. Protein abundance and other physiochemical characteristics do not predict FTR. Therefore, co-analysis of the abundance and synthesis rates of proteins in human muscle is required for correct insight and interpretation.

## Author Contributions

Conceptualization, P.J.L. and J.G.B.; methodology, J.S.B., S.B., J.P., J.A.S., G.L.C. and J.G.B; formal analysis, B.N.S., J.S.B., S.B., C.A.S., J.P. and J.B.L.; investigation, S.O.S., J.A.S., J.B.L., G.L.C. and P.J.L; resources, G.L.C and J.G.B.; data curation, C.A.S and J.G.B. ; writing—original draft preparation, B.N.S., C.A.S. and J.G.B; writing—review and editing, J.S.B. S.B. J.P., S.O.S., J.A.S., J.B.L., G.L.C., and P.J.L.; visualization, B.N.S and C.A.S.; supervision, J.P., S.O.S., J.A.S., J.B.L. and J.G.B.; project administration, J.G.B.; funding acquisition, G.L.C., P.L.J. and J.G.B. All authors have read and agreed to the published version of the manuscript.

## Institutional Review Board Statement

The study was approved by the Liverpool John Moores School of Sport and Exercise Science Research Committee (M18SPS006) and conformed with the Declaration of Helsinki, except registration in a publicly accessible database.

## Informed Consent Statement

Informed consent was obtained from all subjects involved in the study.

## Data Availability Statement

The data presented in this study are available in Suppl Table S1 and the raw mass spectrometry files are deposited in ProteomeXchange, PXD046509.

